# The Endocannabinoid System’s Contribution to Placebo Analgesia

**DOI:** 10.64898/2026.02.25.707676

**Authors:** Rossi Tomin, Kevin Murray, Georgia E. Hadjis, Omar Khalil, Christine Sexton, Stephanie L. Bourke, James S. Khan, David P. Finn, Lauren Y. Atlas, Massieh Moayedi

## Abstract

Placebo analgesia—pain reduction from inert treatments—varies widely across individuals, yet its neurochemical basis remains poorly understood. While endogenous opioids, specifically β-endorphin, contribute to placebo effects, mu-opioid receptor blockade does not fully abolish analgesia, implicating additional systems. Here, we investigate the endogenous cannabinoid (eCB) system’s contribution to placebo analgesia in 48 healthy adults using a validated placebo paradigm with blood sampling. We quantified circulating levels of eCB ligands and β-endorphin at baseline, as well as before and after placebo and control conditions to determine condition-related changes. Individual differences in placebo analgesia were associated with increases in fatty acid amide hydrolase (FAAH) substrates—a composite of anandamide, palmitoylethanolamide, and oleoylethanolamide—but not 2-arachidonoylglycerol or β-endorphin alone. Critically, β-endorphin moderated this relationship: FAAH substrates were strongly associated with pain reduction only when β-endorphin levels were low. These findings provide evidence that eCB and opioid systems interact in a state-dependent manner during placebo analgesia in humans, with implications for understanding individual variability in treatment responses.

## INTRODUCTION

Pain perception is highly modifiable by expectations, beliefs, and the psychosocial context surrounding treatment ^1^. This is most clearly illustrated by placebo analgesia, in which inert treatments reduce pain through psychosocial mechanisms ^2^. Importantly, placebo analgesia is not merely a psychological phenomenon but reflects engagement of endogenous pain modulatory systems ^3-5^. Individuals vary widely in placebo responsiveness ^6-8^, and understanding the biological basis of this variability could enable personalized pain management and improve prediction of treatment outcomes.

The mechanisms underlying placebo analgesia are not unique to placebo and overlap with those engaged by many pain treatments, including active pharmacotherapies ^9^. Understanding the neurobiological basis of placebo effects therefore holds important therapeutic implications, potentially revealing targets to enhance treatment efficacy or develop novel pain management strategies.

Substantial evidence supports a key role for endogenous opioids in mediating placebo analgesia. Administration of the opioid antagonist naloxone attenuates placebo responses ^10-12^, and neuroimaging studies using [11C]-carfentanil reveal placebo-induced changes in µ-opioid receptor availability ^13-15^. A key endogenous ligand of these µ-opioid receptors distributed throughout ascending nociceptive pathways and descending pain modulatory circuits ^16^ is the opioid peptide β-endorphin ^17^. However, naloxone does not completely abolish placebo analgesia ^18-20^, indicating that additional non-opioidergic mechanisms also contribute to these effects ^21,22^.

The endogenous cannabinoid (eCB) system has emerged as a promising candidate for non-opioidergic placebo analgesia ^23^. It comprises cannabinoid receptors (CB_1_R and CB_2_R), endogenous ligands such as arachidonoylethanolamide (anandamide, AEA) and 2-arachidonoylglycerol (2-AG), and metabolizing enzymes, including fatty acid amide hydrolase (FAAH) and monoacylglycerol lipase (MGL). Related signaling molecules, the *N*-acylethanolamines (NAEs) *N*-palmitoylethanolamide (PEA) and *N*-oleoylethanolamide (OEA) also play a role in pain regulation and can increase AEA levels through substrate competition for FAAH ^24-26^. The eCB system is expressed at every level of nociceptive processing and within key descending modulatory circuits. Preclinical studies show that eCB signaling modulates nociception and pain-related behavior, with key sites of action within the descending pain modulatory system ^27,28^. FAAH and MGL inhibitors have antinociceptive effects in multiple animal models of pain ^29-37^. *FAAH* C385A polymorphism, which reduces FAAH activity and is carried by approximately 45% of the population (36.7% heterozygous, 7.8% homozygous)^38,39^, has been associated with lower cold pain sensitivity and reduced post-operative analgesic requirements ^40^. Despite this clear relevance for a role in pain modulation, the role of eCBs in human placebo analgesia remains unclear. While CB_1_R antagonism selectively reduces non-opioidergic placebo responses ^23^, *FAAH* C385A carriers paradoxically exhibit reduced placebo analgesia in one study ^41^, underscoring the complexity of eCB contributions and the need for further investigation.

Critically, converging preclinical evidence indicates that opioid and cannabinoid systems interact. Pretreatment with exogenous cannabinoids enhances opioid analgesia in rodents ^42-45^, and cannabinoid-mediated antinociception can be blocked by µ-opioid receptor antagonists ^8^. However, human studies have yielded inconsistent results ^46,47^, and whether endogenous opioids and eCBs interact to produce placebo analgesia remains unknown, representing a critical gap in our understanding.

In this pre-registered study, we investigated whether circulating eCB ligands contribute to individual differences in placebo analgesia using a validated placebo paradigm ^48^. We measured baseline and condition-evoked changes in circulating levels of eCB system ligands (2-AG and FAAH substrates: AEA, PEA, and OEA) and tested their associations with placebo analgesia. We further examined whether the *FAAH* C385A polymorphism, biological sex, and cannabis use history modulated these relationships. Given evidence for opioid involvement in placebo analgesia and preclinical data suggesting opioid-cannabinoid interactions, we measured β-endorphin levels to test for interactions between the eCB and opioid systems in producing placebo effects in humans.

## RESULTS

### Participants

Participant demographics are provided in Table 1.

**Table 1.** Participant Demographics, *FAAH genotype* and Self-Reported Substance Use.

|  | Male (*n* = 25) | Female (*n* = 23) | Total (*N* = 48) |
| --- | --- | --- | --- |
| Age, mean (SD) | 26.88 (5.80) | 23.08 (4.36) | 25.00 (5.47) |
| **Gender, *n*** |  |  |  |
| Man | 25 | 0 | 25 |
| Woman | 0 | 23 | 23 |
| **Education level, *n*** |  |  |  |
| High school | 6 | 7 | 13 |
| College diploma | 2 | 0 | 2 |
| Bachelor's degree | 13 | 9 | 22 |
| Master's degree | 4 | 5 | 9 |
| Doctoral degree | 0 | 2 | 2 |
| **FAAH genotype, *n*** |  |  |  |
| CC | 12 | 11 | 23 |
| CA | 10 | 12 | 22 |
| AA | 3 | 0 | 3 |
| **Smoking** |  |  |  |
| Current smokers, *n* | 5 | 2 | 7 |
| Packs per week, mean | 0.60 | 1.50 | 1.05 |
| **Alcohol Use** |  |  |  |
| Drinks per week, mean | 1.79 | 1.33 | 1.56 |
| **Cannabis Use, *n*** |  |  |  |
| Current users | 7 | 8 | 15 |
| Previous users | 8 | 2 | 10 |
| Never used | 10 | 13 | 23 |

*N* = 48 participants. SD = standard deviation. Gender categories of transgender, non-binary, and prefer not to specify were offered but not endorsed by any participant. Smoking packs per week are reported among current smokers only.

### Placebo administration reduces pain

Our placebo manipulation successfully reduced pain (Shapiro Wilk’s W = 0.97, p = 0.255; *t*(47) = 4.595, *p* = 3.286 × 10^−5^, Cohen’s *d* = 0.66). Participants reported significantly lower pain ratings in the placebo condition (mean ± SEM = 2.47 ± 0.24) compared to the control condition (mean ± SEM = 3.41 ± 0.18), demonstrating robust placebo analgesia (Figure 1).

**Figure. 1.**
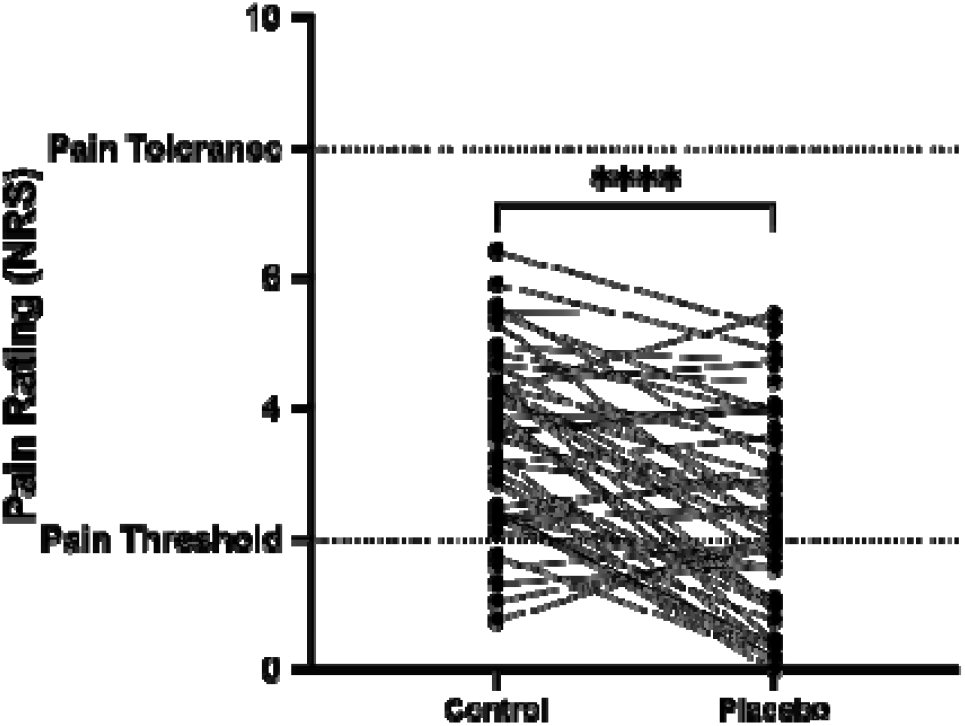
Placebo analgesia. Pain intensity ratings for control and placebo conditions across all participants (N = 48). Individual data points are connected by lines to show within-participant change. Placebo administration significantly reduced pain ratings (p < .0001).

### Endocannabinoid Contributions: FAAH Substrates Are Associated with Placebo Analgesia

We first examined baseline levels of eCB ligands. Linear mixed models revealed no significant associations between placebo analgesia and baseline levels of FAAH substrates or 2-AG (all *p* > .05; Table S1). We therefore examined condition-evoked changes in these ligands— calculated as the difference between post-condition and pre-condition blood samples for each experimental block.

A linear mixed model revealed a significant FAAH substrates-by-condition interaction (β = - 0.697, SE = 0.334, *t(1081*.*0*) = -2.084, *p* = .037; Figure 2 and Table S2), indicating that individuals with greater increases in FAAH substrates experienced stronger placebo analgesia— that is, lower pain ratings in the placebo condition relative to the control condition.

**Figure. 2.**
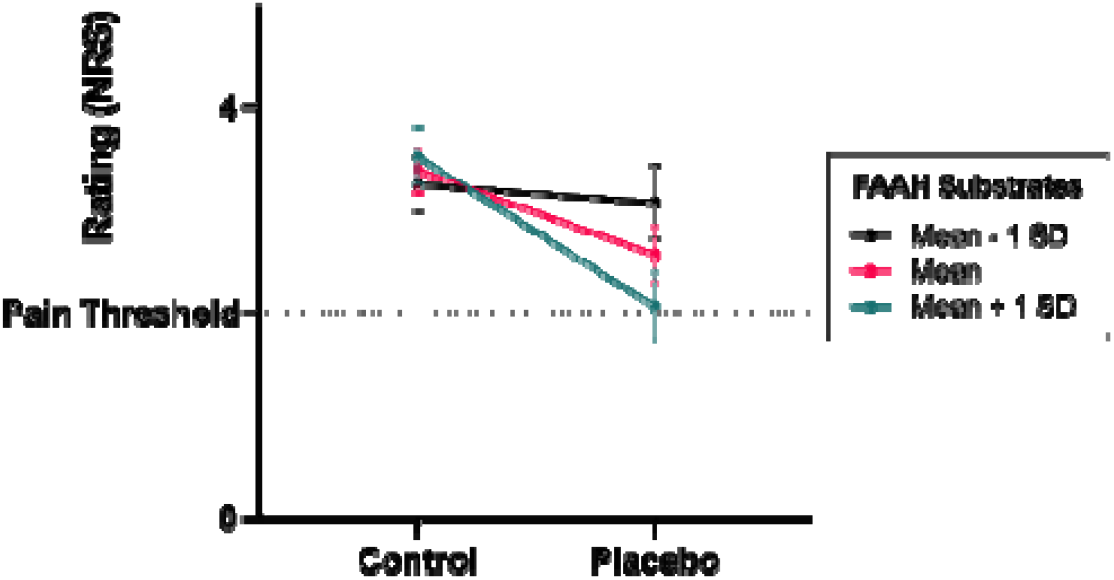
Changes in FAAH substrates are associated with placebo analgesia. Participants with higher changes in FAAH substrate values reported increased placebo analgesia, reflected in lower pain ratings in the placebo condition relative to control. β = -0.697, SE = 0.334, t(1081.0) = -2.084, p = .037; Mean – 1 SD represents a low FAAH substrate level and Mean + 1 SD represents a high FAAH substrate level. Error bars represent ±1 SE.

In contrast, the 2-AG-by-condition interaction was not significant (*p* =.213). Post hoc analyses examining individual FAAH substrates revealed that neither AEA, PEA, nor OEA alone were significantly associated with placebo analgesia (all *p* > .05; Tables S3–S5), indicating that the composite measure captured variance not attributable to any individual analyte alone.

### Opioid Contributions: β-Endorphin Levels Alone Are Not Associated with Placebo Analgesia

We examined whether β-endorphin levels were associated with individual differences in placebo responses. Linear mixed models revealed no significant associations between placebo analgesia and either baseline β-endorphin levels (*p* > .05; Table S6) or condition-evoked changes in β-endorphin (*p* > .05; Table S7).

### β-Endorphin Levels Moderate the FAAH substrates-Placebo Analgesia Relationship

We tested whether β-endorphin and eCB systems interact in human placebo analgesia. A linear mixed model including condition, FAAH substrates, 2-AG, and β-endorphin levels revealed a significant three-way interaction between condition, FAAH substrates, and β-endorphin (β = 1.037, SE = 0.452, *t*(1069.0) = 2.297, *p* = .022; Figure 3; Table S8).

**Figure. 3.**
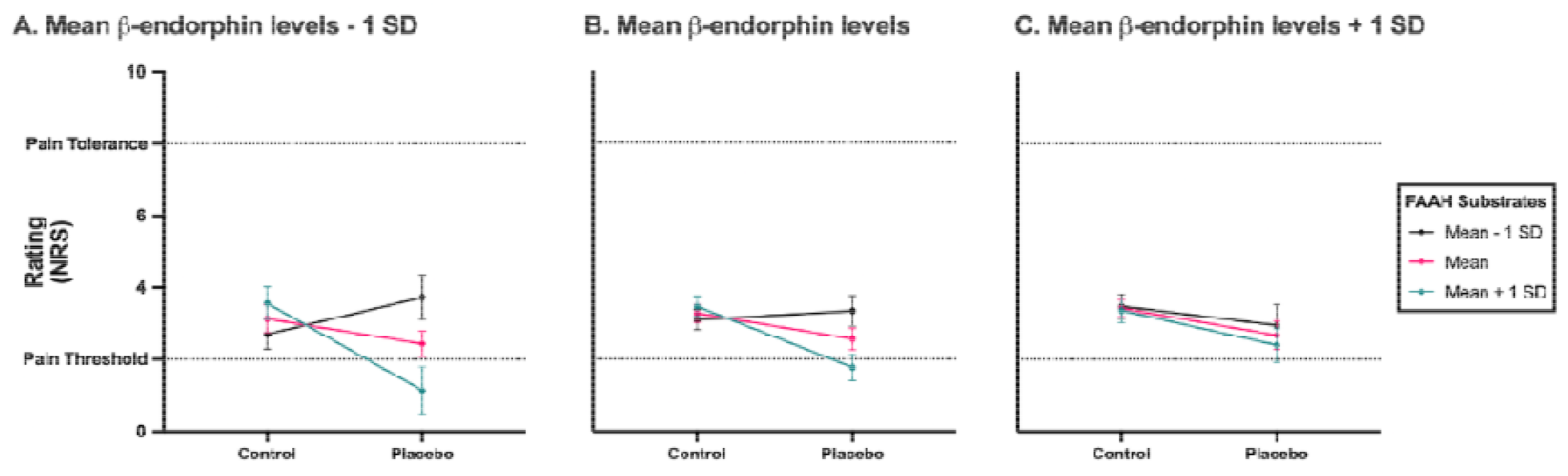
β-Endorphin levels gate endocannabinoid contributions to placebo analgesia. **(A)**□Ratings plotted at low circulating β-endorphin levels (-1 SD). Participants with higher FAAH analyte values (+1 SD) reported increased placebo analgesia, reflected in lower pain ratings in the placebo condition relative to control. (**B)**□Ratings plotted at mean circulating β-endorphin levels. Higher FAAH analyte values (+1 SD) were associated with reduced pain ratings under placebo compared to control. **(C)**□Ratings plotted at high circulating β-endorphin levels (+1 SD). Minimal relationship between FAAH analyte values and placebo analgesia. Error bars represent ±1 SE.

To interpret this interaction, we examined the relationship between FAAH substrates and pain ratings at different levels of β-endorphin using estimated marginal means. When β-endorphin elevations were low (-1 SD), increases in FAAH substrates were strongly associated with lower pain ratings in the placebo condition compared to control (Figure 3A). This association remained evident at mean β-endorphin levels (Figure 3B) but was absent when β-endorphin elevations were high (+1 SD; Figure 3C), indicating that the relationship between FAAH substrates and placebo analgesia differs depending on β-endorphin levels.

### FAAH Genotype, Sex, and Cannabis Use Do Not Moderate Placebo Analgesia

Finally, we examined whether genetic variation in FAAH activity, biological sex, or prior cannabis use modulated placebo responses or their association with circulating eCB levels. We found no significant effects of the *FAAH* C385A polymorphism, cannabis use, or sex on circulating levels of eCBs or related *N*-acylethanolamines (Table S2; all p > .05). Neither FAAH C385A genotype, cannabis use history, nor biological sex significantly moderated placebo analgesia (all p > .05; Table S9).

## DISCUSSION

We provide the first human evidence that eCB and opioid systems interact in a state-dependent manner to produce placebo analgesia in humans. Changes in circulating FAAH substrates (AEA, PEA, OEA) were associated with individual differences in placebo-induced pain reduction, and critically, this relationship was moderated by β-endorphin levels. When β-endorphin levels were minimal, increases in eCB ligands were strongly associated with analgesia; when β-endorphin levels were high, this relationship was no longer evident. These findings reveal that expectation-driven pain relief emerges from coordinated, state-dependent engagement of multiple neuromodulatory systems rather than independent or uniformly additive processes, with important implications for understanding individual variability in placebo responsiveness.

Having established a robust placebo effect, we sought to identify the neurochemical mechanisms underlying individual differences in this response. Importantly, we identified a novel association between placebo analgesia and changes in the eCB system. Increases in circulating FAAH substrates were associated with individual differences in placebo analgesia, providing direct human evidence that eCB mobilization may contribute to expectation-induced pain relief. These findings extend prior pharmacological antagonist studies demonstrating that non-opioidergic mechanisms, including the eCB system, contribute to placebo effects ^23^.

There is good evidence that NAEs, including AEA, OEA, and PEA, have anti-nociceptive and anti-inflammatory properties in preclinical models ^28,49^. While AEA directly engages CB_1_R and CB_2_R, PEA and OEA indirectly modulate eCB signaling through substrate competition at FAAH ^28^. Here, we show that shared variance between the FAAH substrates is associated with placebo analgesia, but the levels of individual analytes are not. It is possible that interactions among NAEs produce the observed effect, rather than any one individual analyte.

From a translational perspective, these results suggest that individuals with higher capacity for eCB mobilization may be particularly responsive to expectation-based interventions. This could inform patient stratification in clinical trials or guide selection of complementary approaches to pharmacotherapy. Interventions that enhance endogenous eCB signaling—such as exercise ^50-53^, dietary omega-3 supplementation ^54^, or stress reduction techniques ^55^—might potentiate placebo responses and improve overall treatment outcomes.

We found that β-endorphin levels alone were not associated with placebo analgesia. This is consistent with prior work showing no placebo-induced changes in plasma β-endorphin when placebo was induced solely through expectation ^56^. While we found no main effect for β-endorphin, our interaction results revealed that β-endorphin facilitate eCB-induced placebo analgesia when mobilization is low.

This gating pattern could reflect several non-mutually exclusive mechanisms within a systems-level framework. One possibility is that robust opioid activation saturates descending pain modulatory circuits, limiting the capacity for additional eCB-mediated modulation. Alternatively, opioid and eCB systems may converge on shared neural substrates within the periaqueductal gray and rostral ventromedial medulla, such that strong engagement of one system reduces the marginal influence of the other. At a molecular level, co-localization ^57^ and heteromerization of µ-opioid and CB_1_Rs ^58^ raise the possibility that activation of one receptor system alters the signaling efficacy of the other. Together, these mechanisms suggest that placebo analgesia emerges from coordinated, state-dependent interactions among neuromodulatory systems, providing a framework for understanding individual differences in expectation-driven pain modulation.

Our findings complement extensive preclinical evidence demonstrating functional interactions between opioid and cannabinoid systems while revealing important distinctions for human placebo analgesia. In rodent models, combined administration of cannabinoids and opioids often produces supra-additive, synergistic antinociceptive effects ^42-44^, and cannabinoid-mediated analgesia can be blocked by µ-opioid receptor antagonists ^59^. However, translation to humans has been inconsistent, with studies reporting variable potentiation, null effects, or even antagonism ^8,46,47^.

We observed no significant effects of *FAAH* C385A genotype or cannabis use history on placebo analgesia. The genotype null finding may reflect limited statistical power to detect effects in the small proportion (∼5%) of AA homozygotes, as our study was not prospectively stratified by genotype. We thus collapsed A-carriers to increase statistical power, but may have reduced the effect by doing so. Regarding cannabis use, the absence of an effect suggests that prior exposure does not fundamentally alter endogenous eCB contributions to placebo analgesia, though future studies with more detailed characterization of use patterns (frequency, recency, THC content) may reveal more nuanced relationships.

Our study has several limitations. First, eCB and β-endorphin dynamics were assessed in peripheral blood, and the relationship between peripheral and central neuromodulator levels remains incompletely understood. Second, our findings are correlational and cannot establish causal relationships between neuromodulator mobilization and analgesia. Third, our sample consisted of healthy volunteers, and whether these mechanisms generalize to chronic pain populations remains to be determined.

These findings suggest several translational opportunities. Circulating eCB and β-endorphin profiles may serve as biomarkers for predicting treatment responses or stratifying participants in clinical trials. Identifying individuals whose analgesia is primarily mediated by eCB versus opioid mechanisms could inform personalized intervention strategies. Future studies combining pharmacological manipulation with multimodal neuroimaging—including PET and fMRI—will be critical for establishing causal mechanisms and mapping the neural circuits through which these systems interact. Extending this work to acute and chronic pain populations will be essential for evaluating clinical relevance.

In summary, we provide the first human evidence that eCB and opioid systems interact to produce placebo analgesia, with β-endorphin levels gating the contribution of eCB signaling to pain relief. These findings fundamentally advance our mechanistic understanding of expectation-driven pain modulation by revealing that multiple neuromodulatory systems operate interdependently rather than independently. By identifying specific neurochemical substrates that explain individual differences in placebo responsiveness, this work opens new avenues for personalized pain management and suggests that optimizing endogenous pain relief may require coordinated engagement of multiple neuromodulatory pathways. This mechanistic insight lays groundwork for developing more sophisticated, biology-based strategies for harnessing the brain’s endogenous analgesic systems in clinical practice.

## METHODS

The study was pre-registered as part of a larger study, on February 25, 2025 on AsPredicted (Wharton Credibility Lab, University of Pennsylvania, PA, USA; https://aspredicted.org/cn83-5zrd.pdf).

### Participants

The study was advertised using posters across the University of Toronto campus and surrounding areas. Seventy participants (35 female and 35 male) aged between 18 and 45 years consented to participate in the study approved by the University of Toronto’s Human Research Ethics Board (Protocol # 45552). To be eligible, participants must not have: (1) a history of opioid use; (2) major neurological or psychiatric diseases; (3) needle phobia; (4) regular prescription medication use; (5) over the counter medication use (in the last 24 hours); (6) history of hypoglycemia, diabetes or pre-diabetes; (7) history of fainting during blood draws;(8) ≥ 16 score on the Beck Depression Inventory (BDI) or endorsed self-harm (> 0 on item (9); ≤ 27 on the Mini Mental State Exam (MMSE); (10) history or presence of chronic pain; (11) visible scarring on the testing site; (12) a heat pain threshold < 36°C; (13) heat pain tolerance > 50°C; (14) coefficient of determination (r^2^) < 0.4 upon completion of calibration procedure (see *Calibration Protocol*, below); (15) suspicion of the placebo manipulation; (16) adverse reactions to blood draws. Twenty-one participants were subsequently excluded from continuing with procedures: 3 for BDI criteria, 10 for blood draw failures, 5 for not being able to follow procedures (e.g., falling asleep during the session), and 3 based on calibration. One participant was excluded from data analysis because their β-endorphin levels were 3 standard deviations outside the mean. This resulted in a final sample of 48 participants (23 female, 25 male, mean ± SD age = 24 ± 5.5 years).

### Experimental Design

Study procedures are presented in Figure 4A. Briefly, after consenting, participants completed two screening questionnaires: the Beck Depression Inventory II and the Mini-Mental State Examination. Those who met inclusion criteria were invited to continue study procedures. First, participants had a cannula inserted in an antecubital vein and a baseline venous blood sample was collected (Sample 1).

**Figure. 4.**
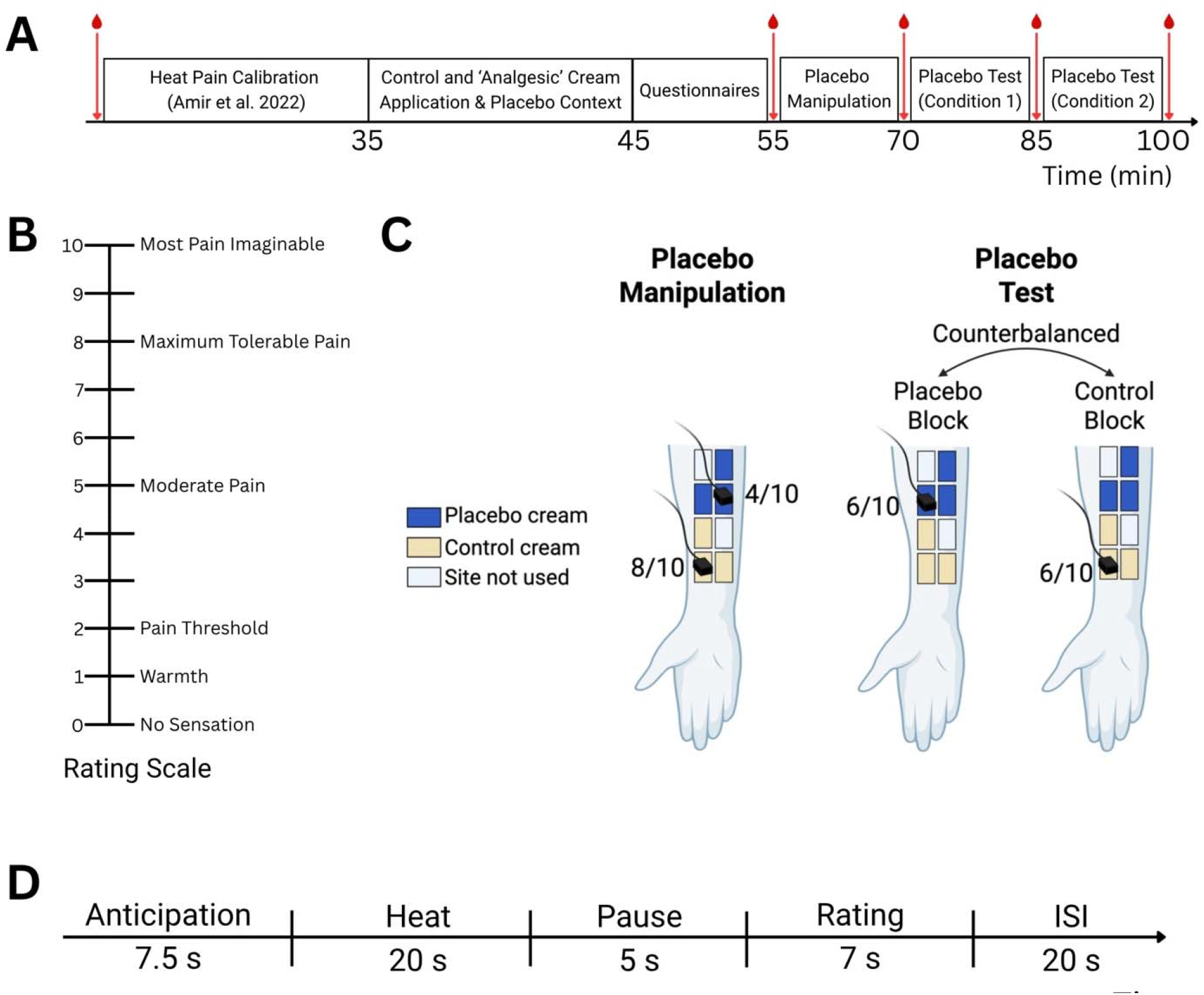
Experimental design. To investigate the contribution of eCB and β-endorphin, and their interaction to placebo analgesia, we used a validated placebo paradigm. **(A)** First, to determine individual stimulus intensities and stimulation sites, we applied an adaptive staircase calibration procedure for thermal heat pain. At the beginning of the placebo paradigm, participants were informed that they would be receiving two creams during the procedure: a highly potent experimental analgesic cream, and a control cream. In reality, both creams provided were Vaseline (petroleum jelly, Vaseline Healing Jelly Original, Unilever). Red lines and drops indicate blood sampling. Next, participants underwent three blocks of testing: **(B)** a placebo manipulation block followed by **(C)** a placebo test, comprising of two conditions blocks: placebo and control, counterbalanced across participants. Blood was drawn before and after each block. In the manipulation block, participants received individually calibrated heat stimuli to elicit a rating of 8/10 to control cream sites, and 4/10 to placebo cream sites. This manipulation served to condition participants and bolster expectations for pain relief. For the next two test blocks, participants received heat stimuli calibrated to elicit a 6/10, either on the control cream sites, or on the ‘analgesic’ cream sites, depending on the block (placebo, control). **(D)** Each block of stimulation consisted of twelve trials. Each trial comprised of a 7.5s anticipation phase where participants saw a red cross hair. This was followed by a 20s heat stimulus. After a 5s pause, participants provided a pain intensity rating, followed by a 20s inter-trial interval. Panel B was made using Biorender.

Participants then completed validated questionnaires related to mood and pain (these are outside the scope of this study, and are not further discussed). Next, they provided another blood sample (Sample 2), underwent placebo manipulation, and then provided another blood sample (Sample 3). Participants then underwent the placebo test phase in two separate blocks, each of which was followed by a blood sample collection (Samples 4 and 5). The order of test blocks (placebo condition first vs. control condition first) was counterbalanced across participants. Blood samples at each timepoint were collected at approximately 15, 70, 90, 110, and 130 minutes from study start, respectively. Sample 2 was collected but not analyzed, as the present study focused on the test phase rather than manipulation-induced changes. Participants then performed other procedures outside the scope of this study. The study session concluded with a debriefing and re-consent process.

### Calibration Protocol

The adapted staircase calibration (ASC) procedure is described in detail elsewhere^60^. Briefly, we delivered noxious heat intensities to 8 sites on the volar forearm (24 total stimuli). Participants provided verbal ratings on a 0 – 10 scale (Figure 4B). We used iterative linear regression to calibrate temperatures to individual pain ratings and select temperatures associated with low (level 2), medium (level 5), or high pain (level 8). Participants with r^2^ > 0.4 had six sites with lowest residual values selected for testing. Detailed methodology is provided in Supplemental Materials.

### Placebo Manipulation and Test Phase

We used a validated placebo paradigm^8,12^ to manipulate and measure placebo analgesia (see Figure 4C). Participants received two inert creams applied to their dominant forearm and were told that one cream (placebo) is a highly potent topical analgesic, and the other is an inert control cream. The placebo cream was extracted from a ‘pharmaceutically labelled’ tube, and was blue, while the control cream was explicitly shown to be Vaseline (Unilever, Toronto, ON, Canada).

In reality, both creams were identical, with the placebo cream being Vaseline with artificial blue colouring.

The placebo manipulation phase (Figure 4C, left panel) consisted of 12 total stimuli (two blocks of six stimuli applied to placebo and control sites, counterbalanced across participants). The placebo site received a lower intensity heat stimulus (4 on the 10-point scale) while the control site received a higher intensity heat stimulus (8 on the 10-point scale) to reinforce participants’ beliefs that the analgesic cream was active and potent.

During the test phase (Figure 4C, right panel), 24 identical heat stimuli (calibrated to level 6 on the 10-point scale) were given to a new set of placebo and control sites, with order counterbalanced across participants. The test phase consisted of 12 heat trials administered first to the control sites, or first to the placebo sites, followed by 12 heat stimuli administered to sites from the condition. The trial structure is shown in Figure 4D. Briefly, each trial consisted of a 7.5-s anticipation phase (red crosshair on screen), followed by the 20-s noxious heat stimulus, a short pause and a pain rating screen, where participants reported their pain intensity rating using the same scale as the ASC (see Figure 4B).

### Sample Collection

Study sessions were scheduled between 7:00 am and 10:30 am to account for circadian fluctuations in eCBs ^61-64^. In addition, participants were required to fast—this included all food and drinks except for water for 10 hours and all over the counter medications for 24 hours—prior to the testing session. A standardized snack was provided to participants at the beginning of the session (MadeGood Mornings, Riverside Natural Foods, Toronto, Canada). Throughout the session, 5 blood samples were taken by using a permanent venous catheter (BD Cathena− Safety IV Catheter with Wings BD Multiguard− Technology with wings 22GA X 1.00IN). Samples were collected into BD Vacutainer® Plus Plastic Serum Tubes at approximately 15, 70, 90, 110, and 130 minutes from study start (Samples 1–5, respectively). The first blood draw also included a sample of whole blood collected into a BD Vacutainer® spray coated K2EDTA blood tube for genetic analysis.

#### Sample Processing

Blood was preprocessed at the University of Toronto’s Faculty of Dentistry. Samples were left at room temperature for 30–60 minutes to allow clotting. Samples were then spun at 1400 g for 15 minutes at 4°C. Serum was then aliquoted into 2 separate 1.5 mL microfuge tubes (Sarstedt).

Samples were stored at -80°C, until they were shipped in a single batch to David P. Finn’s laboratory at the University of Galway for analysis.

#### Quantification of β-Endorphin from Human Serum via ELISA

Serum concentrations of β-endorphin were analysed by competitive enzyme immunoassay using a human β-endorphin ELISA Kit (HUFI00498, Assay Genie, Dublin, Ireland), as per manufacturer instructions (see Supplementary Materials for detailed methods).

#### Lipid Extraction of Endocannabinoids and Related NAEs in Serum

Serum levels of 2-AG, AEA, PEA and OEA were analysed as previously described^65,66^. Briefly, 200 μL of serum from each sample was spiked with 20 μL of 100% acetonitrile (ACN) containing the eCB and NAE internal standard mix (50 ng of 2-AG-d8, 2.5 ng of AEA-d8, 2.5 ng of OEA-d4 and 2.5 ng of PEA-d4). Proteins were precipitated with 1mL of 100% ACN containing 0.1% formic acid. The precipitated proteins were pelleted by centrifugation at 18,620 g for 15 minutes at 4°C. The supernatant was then filtered, and the collected eluate was vortexed, and 500 μL of the total volume was dried down and reconstituted in 20 μL and transferred into a HPLC vial (see Supplementary Materials for detailed methods and LC-MS/MS assay parameters).

#### FAAH SNP (C385A) Genotyping

Genomic DNA (gDNA) was extracted using standard protocols (Nucleospin® DNA isolation kit, Macherey-Nagel, Düren, Germany). All samples were genotyped for the C385A *FAAH* SNP (rs324420) using predesigned TaqMan primers and universal genotyping master mix (Life Technologies, Ireland). Genotyping assays were performed according to manufacturers’ protocols using StepOne Plus PCR machine (Applied Biosystems, Ireland) and allelic discrimination software (Applied Biosystems).

### Statistical Analyses

We pre-registered (AsPredicted #: 214,720; https://aspredicted.org/cn83-5zrd.pdf) the hypothesis that individuals exhibiting greater placebo analgesia (defined as control pain intensity minus placebo pain intensity) would show increases in circulating eCBs, and that this relationship would be moderated by *FAAH* C385A genotype and sex. All primary dependent variables, predictors (changes in eCB levels, *FAAH* genotype, sex, and relevant covariates), and modeling strategies (linear mixed models) were preregistered. Given the well-established role of endogenous opioids in placebo analgesia^11-13^ and preclinical evidence demonstrating functional interactions between the eCB and opioidergic systems^8,42-44^, we extended our analyses to explore β-endorphin contributions.

We performed linear mixed models using the R package *GAMLj3*^*67-69*^ implemented in jamovi (version 2.7). Models were estimated using restricted maximum likelihood (REML). Given the repeated measures collected in this study, we specified a first-order autoregressive (AR(1)) residual covariance structure account for temporal autocorrelation between adjacent trials — an assumption that is more appropriate than independence for serially collected repeated measures. Confidence intervals for fixed-effect coefficients were estimated using parametric bootstrapping (1,000 replications), which provides robust interval estimates with moderate sample sizes (N = 48) without relying on asymptotic distributional assumptions. Degrees of freedom were estimated using the Kenward-Roger approximation, which applies a small-sample correction to the denominator degrees of freedom and the covariance matrix of fixed effects, yielding more accurate F-tests in designs with limited clusters than the Satterthwaite method. Statistical significance was set at α = .05 for all tests. No corrections for multiple comparisons were applied, as primary hypotheses were specified *a priori* in the preregistration.

We performed five linear mixed models with the same structure as described above. Specifically, two models investigated the contribution of baseline eCB/NAE and β-endorphin to placebo analgesia, respectively. Two further models investigated the contribution of placebo-induced changes in eCB/NAE, β-endorphin to placebo analgesia, respectively. Next, given that our primary analyses revealed significant associations between FAAH substrates (see *Data Reduction*, below) and placebo analgesia we tested an integrated model including both β-endorphin levels and eCBs to examine potential interactions between these systems in producing placebo effects.

Finally, three post-hoc models were performed to parse the contribution of each FAAH substrate—AEA, OEA, PEA. Each model had a random slope and intercept with participant ID and condition as random effects.

#### Calculation of Condition-Evoked Changes

For each participant, condition-evoked changes in eCB ligands and β-endorphin were calculated separately for each experimental condition (placebo and control), calculated as the difference between post-condition and pre-condition blood samples for each experimental block. Because test blocks were counterbalanced, the specific blood samples used for each calculation depended on block order:

- For the **first test block** (either placebo or control): Change = Sample 4 - Sample 3
- For the **second test block** (either control or placebo): Change = Sample 5 - Sample 4

This approach ensured that each condition’s change score was calculated relative to the blood sample collected immediately before that specific test block, accounting for any carryover effects and providing condition-specific estimates of neuromodulator mobilization.

#### Data Reduction

To reduce the dimensionality of collinear predictors, principal component analyses (PCAs) were performed on baseline levels and condition-evoked changes in AEA, OEA and PEA. PCAs were performed in IBM SPSS Statistics for Mac (Version 28). The data’s suitability for reduction was assessed using Bartlett’s test of sphericity (*p* < .05), and sampling adequacy was confirmed via the Kaiser-Meyer-Olkin test (threshold > .5). Predictors that did not satisfy these criteria were excluded from their respective PCAs and incorporated into subsequent analyses as individual variables. Components were selected based on an eigenvalue threshold > 1. In cases where more than one component was extracted, an orthogonal varimax rotation algorithm was applied to ensure the resulting components were orthogonal, thereby minimizing collinearity in downstream analyses. The PCA yielded a single component termed ‘FAAH substrates’ (see Tables S10 – S13 for assumption tests and loadings).

#### Placebo Analgesia Determination

We first examined whether the placebo manipulation produced analgesia. We first tested that data met normality assumptions with the Shapiro-Wilk’s test. Next, a two-sided paired-samples *t*-test comparing pain ratings between placebo and control conditions. The effect size (Cohen’s d) was computed in jamovi.

#### Baseline eCB and β-Endorphin Levels

We examined whether baseline levels (Sample 1) of FAAH substrates, 2-AG, or β-endorphin were associated with placebo analgesia using separate linear mixed models for eCBs and β-endorphin. These models included condition (placebo vs. control), and the baseline measure as fixed effects, along with all two-way and three-way interactions. Covariates included block order (placebo-first vs. control-first), *FAAH* C385A genotype (A-carrier vs. non-carrier), cannabis use (never used vs. previously or currently using), sex, and testing time (to account for circadian effects). Participant ID was modeled as a random effect with random slopes for condition.

#### Changes in Circulating eCBs and Placebo Analgesia

To investigate how condition-evoked changes in eCBs affected pain ratings, we utilized linear mixed models with random slopes and intercepts. The primary model included condition (placebo vs. control), FAAH substrates, and 2-AG as fixed effects, along with all two-way and three-way interactions, with the same covariates and random effects structure described above. Post hoc analyses examined individual FAAH substrates (AEA, OEA, PEA) separately to determine whether any single analyte drove the composite effect.

#### Changes in Circulating β-endorphin and Placebo Analgesia

We assessed associations with changes in β-endorphin levels using linear mixed models parallel to those for eCBs. Models included condition, change scores for β-endorphin levels, and their interaction as fixed effects, with the same covariates described above.

#### Interaction Between eCB and β-endorphin in Placebo Analgesia

To test whether eCB and opioid systems interact to produce placebo analgesia, we fit a linear mixed model including condition, FAAH substrates, 2-AG, and β-endorphin change scores as predictors, along with all two-way and three-way interactions. The critical test was the three-way interaction between condition, FAAH substrates, and β-endorphin. To interpret significant interactions, we calculated estimated marginal means and examined simple slopes at low (−1 SD), mean, and high (+1 SD) levels of β-endorphin.

## Funding

This work was supported by a Canadian Institutes of Health Research Project Grant PJT183703.

LYA’s contribution to this research was supported in part by the Intramural Research Program of the National Institutes of Health (NIH). The contributions of the NIH author are considered Works of the United States Government. The findings and conclusions presented in this paper are those of the author and do not necessarily reflect the views of the NIH or the U.S. Department of Health and Human Services.

MM holds a Canada Research Chair (Tier 2) in Pain NeuroImaging and is supported by a University of Toronto Centre for the Study of Pain Scientist Award. RT was supported by an Ontario Graduate Scholarship, and the Wilson G. Harron Trust at the University of Toronto’s Faculty of Dentistry.

These data have been published as a preprint on bioRxiv: https://www.biorxiv.org/content/10.64898/2026.02.25.707676v1

These data were presented at the ACNP meeting in 2026.

## Author contributions

Conceptualization: MM, LYA, DPF Methodology: JSK, RT, SLB, LYA, MM Scripting: OK

Data Collection: RT, CS Analyte Analysis: KM

Statistical Analysis: RT, LYA, MM, GEH Supervision: MM, LYA, DPF Writing—original draft: RT, MM, LYA

Writing—review & editing: KM, GEH, OK, CS, SLB, JSK, DPF, LYA

## Competing interests

Authors declare that they have no competing interests.

## Data and materials availability

All relevant data is appended.

## Notes

### Competing Interest Statement

The authors have declared no competing interest.

### Summary of Updates

Linear Mixed Models updated and reran. Updated all figures and tables.

## REFERENCES

1. Benedetti, F. Placebo effects: Understanding the other side of medical care, (Oxford University Press, 2021).

2. Pagnini, F., et al. Enacting the mind/body connection: the role of self-induced placebo mechanisms. Humanities and Social Sciences Communications 11, 977 (2024).

3. Atlas, L.Y., et al. Dissociable influences of opiates and expectations on pain. J Neurosci 32, 8053–8064 (2012).

4. Bingel, U., et al. The effect of treatment expectation on drug efficacy: imaging the analgesic benefit of the opioid remifentanil. Sci Transl Med 3, 70ra14 (2011).

5. Schenk, L.A., Sprenger, C., Geuter, S. & Buchel, C. Expectation requires treatment to boost pain relief: an fMRI study. Pain 155, 150–157 (2014).

6. Crawford, L.S., et al. Stimulus-independent and stimulus-dependent neural networks underpin placebo analgesia responsiveness in humans. Commun Biol 6, 569 (2023).

7. Wang, Y., Chan, E., Dorsey, S.G., Campbell, C.M. & Colloca, L. Who are the placebo responders? A cross-sectional cohort study for psychological determinants. Pain 163, 1078–1090 (2022).

8. Wager, T.D., Atlas, L.Y., Leotti, L.A. & Rilling, J.K. Predicting individual differences in placebo analgesia: contributions of brain activity during anticipation and pain experience. J Neurosci 31, 439–452 (2011).

9. Hoffman, G.A., Harrington, A. & Fields, H.L. Pain and the placebo: what we have learned. Perspect Biol Med 48, 248–265 (2005).

10. Levine, J.D., Gordon, N.C. & Fields, H.L. The mechanism of placebo analgesia. Lancet 2, 654–657 (1978).

11. Benedetti, F. The opposite effects of the opiate antagonist naloxone and the cholecystokinin antagonist proglumide on placebo analgesia. Pain 64, 535–543 (1996).

12. Eippert, F., et al. Activation of the opioidergic descending pain control system underlies placebo analgesia. Neuron 63, 533–543 (2009).

13. Wager, T.D., Scott, D.J. & Zubieta, J.K. Placebo effects on human mu-opioid activity during pain. Proc Natl Acad Sci U S A 104, 11056–11061 (2007).

14. Scott, D.J., et al. Placebo and nocebo effects are defined by opposite opioid and dopaminergic responses. Arch Gen Psychiatry 65, 220–231 (2008).

15. Zubieta, J.K., et al. Placebo effects mediated by endogenous opioid activity on mu-opioid receptors. Journal of Neuroscience 25, 7754–7762 (2005).

16. Stein, C. The control of pain in peripheral tissue by opioids. N Engl J Med 332, 1685–1690 (1995).

17. Zhang, R.R., Zhang, W.C., Wang, J.Y. & Guo, J.Y. The opioid placebo analgesia is mediated exclusively through mu-opioid receptor in rat. Int J Neuropsychopharmacol 16, 849–856 (2013).

18. Benedetti, F., Arduino, C. & Amanzio, M. Somatotopic activation of opioid systems by target-directed expectations of analgesia. J Neurosci 19, 3639–3648 (1999).

19. Benedetti, F. & Amanzio, M. The neurobiology of placebo analgesia: from endogenous opioids to cholecystokinin. Prog Neurobiol 52, 109–125 (1997).

20. Gracely, R.H., Dubner, R., Wolskee, P.J. & Deeter, W.R. Placebo and naloxone can alter post-surgical pain by separate mechanisms. Nature 306, 264–265 (1983).

21. Scott, D.J., et al. Individual differences in reward responding explain placebo-induced expectations and effects. Neuron 55, 325–336 (2007).

22. Pecina, M. & Zubieta, J.K. Molecular mechanisms of placebo responses in humans. Mol Psychiatry 20, 416–423 (2015).

23. Benedetti, F., Amanzio, M., Rosato, R. & Blanchard, C. Nonopioid placebo analgesia is mediated by CB1 cannabinoid receptors. Nat Med 17, 1228–1230 (2011).

24. Starowicz, K. & Finn, D.P. Cannabinoids and Pain: Sites and Mechanisms of Action. Adv Pharmacol 80, 437–475 (2017).

25. Corcoran, L., Roche, M. & Finn, D.P. The Role of the Brain’s Endocannabinoid System in Pain and Its Modulation by Stress. Int Rev Neurobiol 125, 203–255 (2015).

26. Di Marzo, V., Bifulco, M. & De Petrocellis, L. The endocannabinoid system and its therapeutic exploitation. Nat Rev Drug Discov 3, 771–784 (2004).

27. Finn, D.P., et al. Cannabinoids, the endocannabinoid system, and pain: a review of preclinical studies. Pain 162, S5–S25 (2021).

28. Woodhams, S.G., Chapman, V., Finn, D.P., Hohmann, A.G. & Neugebauer, V. The cannabinoid system and pain. Neuropharmacology 124, 105–120 (2017).

29. Booker, L., et al. The fatty acid amide hydrolase (FAAH) inhibitor PF-3845 acts in the nervous system to reverse LPS-induced tactile allodynia in mice. Br J Pharmacol 165, 2485–2496 (2012).

30. Kinsey, S.G., Naidu, P.S., Cravatt, B.F., Dudley, D.T. & Lichtman, A.H. Fatty acid amide hydrolase blockade attenuates the development of collagen-induced arthritis and related thermal hyperalgesia in mice. Pharmacol Biochem Behav 99, 718–725 (2011).

31. Kinsey, S.G., Long, J.Z., Cravatt, B.F. & Lichtman, A.H. Fatty acid amide hydrolase and monoacylglycerol lipase inhibitors produce anti-allodynic effects in mice through distinct cannabinoid receptor mechanisms. J Pain 11, 1420–1428 (2010).

32. Sagar, D.R., Jhaveri, M. & Chapman, V. Targeting the cannabinoid system to produce analgesia. Curr Top Behav Neurosci 1, 275–287 (2009).

33. Jhaveri, M.D., Richardson, D., Kendall, D.A., Barrett, D.A. & Chapman, V. Analgesic effects of fatty acid amide hydrolase inhibition in a rat model of neuropathic pain. J Neurosci 26, 13318–13327 (2006).

34. Guindon, J., Guijarro, A., Piomelli, D. & Hohmann, A.G. Peripheral antinociceptive effects of inhibitors of monoacylglycerol lipase in a rat model of inflammatory pain. Br J Pharmacol 163, 1464–1478 (2011).

35. Thomas, A., Okine, B.N., Finn, D.P. & Masocha, W. Peripheral deficiency and antiallodynic effects of 2-arachidonoyl glycerol in a mouse model of paclitaxel-induced neuropathic pain. Biomed Pharmacother 129, 110456 (2020).

36. Diester, C.M., Lichtman, A.H. & Negus, S.S. Behavioral Battery for Testing Candidate Analgesics in Mice. II. Effects of Endocannabinoid Catabolic Enzyme Inhibitors and ?9-Tetrahydrocannabinol. J Pharmacol Exp Ther 377, 242–253 (2021).

37. Grim, T.W., et al. Combined inhibition of FAAH and COX produces enhanced anti-allodynic effects in mouse neuropathic and inflammatory pain models. Pharmacol Biochem Behav 124, 405–411 (2014).

38. Ensembl Database. rs324420 SNP: Population Genetics. Vol. 2025 (EMBL’s European Bioinformatics Institute, Online, 2025).

39. Sipe, J.C., Waalen, J., Gerber, A. & Beutler, E. Overweight and obesity associated with a missense polymorphism in fatty acid amide hydrolase (FAAH). Int J Obes (Lond) 29, 755–759 (2005).

40. Cajanus, K., et al. Effect of endocannabinoid degradation on pain: role of FAAH polymorphisms in experimental and postoperative pain in women treated for breast cancer. Pain 157, 361–369 (2016).

41. Pecina, M., et al. FAAH selectively influences placebo effects. Mol Psychiatry 19, 385–391 (2014).

42. Cichewicz, D.L. Synergistic interactions between cannabinoid and opioid analgesics. Life Sci 74, 1317–1324 (2004).

43. Maguire, D.R., Yang, W. & France, C.P. Interactions between mu-opioid receptor agonists and cannabinoid receptor agonists in rhesus monkeys: antinociception, drug discrimination, and drug self-administration. J Pharmacol Exp Ther 345, 354–362 (2013).

44. Welch, S.P. Interaction of the cannabinoid and opioid systems in the modulation of nociception. Int Rev Psychiatry 21, 143–151 (2009).

45. Rodriguez-Munoz, M., Onetti, Y., Cortes-Montero, E., Garzon, J. & Sanchez-Blazquez, P. Cannabidiol enhances morphine antinociception, diminishes NMDA-mediated seizures and reduces stroke damage via the sigma 1 receptor. Mol Brain 11, 51 (2018).

46. Naef, M., et al. The analgesic effect of oral delta-9-tetrahydrocannabinol (THC), morphine, and a THC-morphine combination in healthy subjects under experimental pain conditions. Pain 105, 79–88 (2003).

47. Roberts, J.D., Gennings, C. & Shih, M. Synergistic affective analgesic interaction between delta-9-tetrahydrocannabinol and morphine. Eur J Pharmacol 530, 54–58 (2006).

48. Wager, T.D., et al. Placebo-induced changes in FMRI in the anticipation and experience of pain. Science 303, 1162–1167 (2004).

49. Paulsen, R.T. & Burrell, B.D. Comparative studies of endocannabinoid modulation of pain. Philos Trans R Soc Lond B Biol Sci 374, 20190279 (2019).

50. You, T., Disanzo, B.L., Wang, X., Yang, R. & Gong, D. Adipose tissue endocannabinoid system gene expression: depot differences and effects of diet and exercise. Lipids Health Dis 10, 194 (2011).

51. Brellenthin, A.G., Crombie, K.M., Hillard, C.J. & Koltyn, K.F. Endocannabinoid and Mood Responses to Exercise in Adults with Varying Activity Levels. Med Sci Sports Exerc 49, 1688–1696 (2017).

52. Feuerecker, M., et al. Effects of exercise stress on the endocannabinoid system in humans under field conditions. Eur J Appl Physiol 112, 2777–2781 (2012).

53. Sparling, P.B., Giuffrida, A., Piomelli, D., Rosskopf, L. & Dietrich, A. Exercise activates the endocannabinoid system. Neuroreport 14, 2209–2211 (2003).

54. Mandal, G., et al. The dietary ligands, omega-3 fatty acid endocannabinoids and short-chain fatty acids prevent cytokine-induced reduction of human hippocampal neurogenesis and alter the expression of genes involved in neuroinflammation and neuroplasticity. Mol Psychiatry 30, 5338–5355 (2025).

55. Bluett, R.J., et al. Endocannabinoid signalling modulates susceptibility to traumatic stress exposure. Nature communications 8, 14782 (2017).

56. Roelofs, J., et al. Expectations of analgesia do not affect spinal nociceptive R-III reflex activity: an experimental study into the mechanism of placebo-induced analgesia. Pain 89, 75–80 (2000).

57. Salio, C., et al. CB1-cannabinoid and mu-opioid receptor co-localization on postsynaptic target in the rat dorsal horn. Neuroreport 12, 3689–3692 (2001).

58. Hojo, M., et al. mu-Opioid receptor forms a functional heterodimer with cannabinoid CB1 receptor: electrophysiological and FRET assay analysis. J Pharmacol Sci 108, 308–319 (2008).

59. Haller, V.L., Stevens, D.L. & Welch, S.P. Modulation of opioids via protection of anandamide degradation by fatty acid amide hydrolase. Eur J Pharmacol 600, 50–58 (2008).

60. Amir, C., et al. Test-Retest Reliability of an Adaptive Thermal Pain Calibration Procedure in Healthy Volunteers. J Pain 23, 1543–1555 (2022).

61. Hanlon, E.C. Impact of circadian rhythmicity and sleep restriction on circulating endocannabinoid (eCB) N-arachidonoylethanolamine (anandamide). Psychoneuroendocrinology 111, 104471 (2020).

62. Hanlon, E.C., et al. Circadian rhythm of circulating levels of the endocannabinoid 2-arachidonoylglycerol. J Clin Endocrinol Metab 100, 220–226 (2015).

63. Kesner, A.J. & Lovinger, D.M. Cannabinoids, Endocannabinoids and Sleep. Front Mol Neurosci 13, 125 (2020).

64. Vaughn, L.K., et al. Endocannabinoid signalling: has it got rhythm? Br J Pharmacol 160, 530–543 (2010).

65. Fatemi, S.A., et al. The Contribution of Baseline Circulating Endocannabinoids to Individual Differences in Human Pain Sensitivity: A Quantitative Sensory Testing Study. bioRxiv (2025).

66. Bourke, S.L., et al. Clinical measures in chronic neuropathic pain are related to the Kennedy and endocannabinoid pathways. Eur J Clin Invest 55, e14351 (2025).

67. The jamovi project. jamovi. (2024).

68. R Core Team. R: A Language and Environment for Statistical Computing. (R Foundation for Statistical Computing, Vienna, Austria, 2024).

69. Gallucci, M. GAMLj: General analysis for linear models. (2019).

