## Supplementary Materials for "The Endocannabinoid System’s Contribution to Placebo Analgesia"

Rossi Tomin *et al.*

**This PDF file includes:**

Supplementary Materials

Supplementary Results

Figure S1

Tables S9-S13, S15

Supplementary References

Supplementary Materials

*Calibration Protocol*

Our calibration protocol consisted of an ascending staircase calibration (ASC) procedure (1). This method has been validated to identify individual pain threshold and tolerance values, and the reliability of the ratings to given temperatures within participants. ASC involves the application of three noxious heat intensities (low, medium, high) to eight skin sites on the dominant volar forearm using a 30 mm x 30 mm thermal probe (T11 probe, TCS II, QSTLab, Strasbourg, France), with temperatures determined through an iterative regression procedure. All stimuli are 20 s in duration, including a 1.5 s ramp-up and 1.5 s ramp-down between the baseline and target temperature (adjusted for each target temperature)—i.e., the stimulus was at target temperature plateau for 17 s. After each heat stimulus, participants provide verbal ratings on a 10-point scale (0 = no sensation, 1 = warmth, 2 = pain threshold, 5 = moderate pain, 8 = maximum tolerable pain, 10 = most pain imaginable). Stimuli are applied to sites 1 through 8, repeated three times, for a total of 24 stimuli. Intensities were pseudorandomized to avoid habituation and sensitization at the sites. The first three temperatures applied are always 41°C, 44°C and 47°C, to establish initial ratings to low, medium and high intensity levels. A linear regression uses these ratings to predict the temperature to be applied on the subsequent site based on the intensity (low, medium, or high) designated for that next site. The rating corresponding to the 4th site is then added to the regression to predict the following temperature to be used on the 5th site, and this is repeated in an iterative fashion for 24 trials (three per site, eight sites). Upon completion of all trials, the r^2^ value from the correlation between temperatures and ratings was used to determine eligibility to continue to the placebo paradigm (r^2^ > 0.4; 3 participants deemed ineligible). For the participants with r^2^ > 0.4, the skin six sites with the lowest residual values were selected for cream application and placebo paradigm testing (three sites for placebo cream application, three sites for control cream).

*Quantification of β-Endorphin from Human Serum via ELISA*

Serum concentrations of β-endorphin were analysed by competitive enzyme immunoassay using a human β-endorphin ELISA Kit (HUFI00498, Assay Genie, Dublin, Ireland), as per manufacturer instructions. A range-finding assay was performed ahead of sample testing to determine the optimal dilution factor, whereby samples would fall in the linear portion of the 4-parameter logistic curve. This kit has a sensitivity of 9.375 pg/mL, and a range of 15.625-1000 pg/mL. Samples were removed from the -80°C freezer and left to thaw at room temperature for two hours. All reagents and the standard curve were prepared as per manufacturer’s protocol. Samples were diluted with sample dilution buffer. The pre-coated 96-well plate was washed twice with wash buffer using a plate washer (Agilent BioTek ELx50, Dublin, Ireland). 50µl of the standards and samples were pipetted into the plate’s wells in duplicate, followed immediately with 50µl of Biotin-labelled antibody working solution into each well. The plate was then sealed and incubated at 37°C for 45 minutes. The plate was removed and washed three more times with wash buffer. 100µl of HRP-Streptavidin conjugate was added to each well, before the plate was covered and incubated at 37°C for 30 minutes. Following this, five further washes were completed. 90µl of TMB substrate was added to each well, then the plate was covered and incubated in the dark for 10 minutes, allowing a blue colour to develop. As soon as this colour developed, the plate was removed from the incubator and 50µl of stop solution was added to each well and mixed thoroughly, leading to a yellow colour change. The plate was again covered and immediately brought to the microplate reader (HIDEX Sense Microplate Reader, Tulku, Finland), where the absorbance was read at 450nm. A four-parameter logistic curve was then plotted to establish the concentration of the samples relative to the optical density (OD) of the known standard curve.

*Lipid Extraction of eCB and Related NAEs in Serum*

Prior to lipid extraction, frozen serum was placed on ice for 60 minutes to defrost the samples while minimizing enzymatic activity affecting the eCBs and NAEs in the samples. Samples were then centrifuged at 18,620 g for 15 minutes at 4°C (Hettich® centrifuge Mikro 22R, Hettich, Germany) to ensure homogeneity of endocannabinoid and NAE levels throughout, before transferring a total volume of 200μL of serum into a 1.5mL microfuge tube (SafeSeal 1.5mL, #72.706, Sarstedt) for the extraction process. Each sample was spiked with 20μL of 100% acetonitrile (ACN) containing the eCB and NAE internal standard mix (50 ng of 2-AG-d8, 2.5 ng of AEA- d8, 2.5 ng of OEA-d4 and 2.5 ng of PEA-d4). Samples were then vortexed and allowed to equilibrate for 10 minutes on ice. For protein precipitation, 1ml of 100% ACN containing 0.1% formic acid, maintained at 4°C, was added to each sample and immediately vortexed and incubated on ice for 30 minutes. The precipitated proteins were pelleted by centrifugation at 18,620 g for 15 minutes at 4°C. A filter (Fisherbrand^TM^ Non-sterile PTFE Hydrophyl, 25 mm, 0.45 μM Syringe Filter, Fisher Scientific, Ireland) was placed in a 5mL SafeSeal tube (SARSTEDT, Ireland) for each sample. 1 mL syringes were attached to the filters by carefully inserting the syringe neck into the filter inlet. The plunger/piston from each 1ml syringe was removed and saved in a sterile container. 1100 μL of the supernatant was removed without disturbing the pellet and loaded into the syringe. The supernatant was allowed to flow through the filter until the syringe was emptied. Once the supernatant had flowed through the syringe into the microcentrifuge tube, 700 μL of 100% ACN was added to the syringe to displace the dead volume of the filter. The syringe plunger was slowly re-inserted, and the 100% ACN was pushed through the filter. The collected eluate was vortexed, and 500μL of the total volume was transferred to a new 1.5 mL microcentrifuge tube and dried down at 45°C for approximately 2 hours in a centrifugal concentrator (Eppendorf Concentrator Plus Complete System, Davidson and Hardy Ltd, Ireland). Samples were then re-constituted in 100μL of 100% ACN, vortexed and dried down again at 45°C for approximately 30 minutes. Each sample was reconstituted in 20μL of 100% ACN before transferring it to a HPLC vial.

*Quantification of eCB and Related NAEs in Serum by LC-MS/MS*

A standard curve using a 1/4 serial dilution was prepared by adding 25 μL of 100% ACN containing a known fixed amount of non-deuterated internal standard (125 ng 2-AG and 12.5 ng AEA, PEA, OEA and) to 75 μL of 100% ACN into the highest standard (Standard 10). 20 μL of 100% CAN containing a known fixed amount of deuterated internal standard was added to each standard, using the same stock used in the samples. Standards were dried down at 45°C for approximately 30 min, then re- constituted in 100% ACN, vortexed and dried down a second time at 45°C for 30 min. Each standard was reconstituted in 20 μL of 100% ACN before transferring it to a HPLC vial. Mobile phases comprised of Solution A (HPLC-grade water with 0.1% (v/v) formic acid) and Solution B (100% acetonitrile with 0.1% (v/v) formic acid) with a flow rate of 0.2 ml/min, onto a Waters Atlantis T3 HPLC column (3 μm particle size dimension, 100 mm length, 2.1 mm diameter; Waters, UK). Reversed-phase gradient elution was initiated at 45% solution B for 1 minute, then ramped linearly up to 100% solution B for 4 minutes and held at 100% solution B until 12 minutes. At 12.1 minutes, when the assay run finished, the gradient returned to initial conditions for a further 4-5 minutes to re- equilibrate the column before the next injection. Analyte detection was carried out in electrospray-positive ionisation mode on an Agilent 1260 Infinity II HPLC system coupled to a SCIEX QTRAP 4500 mass spectrometer operated in triple quadrupole mode (SCIEX Ltd, Phoenix House Lakeside Drive Centre Park, United Kingdom). Ratiometric quantification was calculated using Skyline (MacCoss Lab Software, University of Washington, Seattle). eCB and NAE concentrations in unknown samples was calculated from the analyte/internal standard peak area response ratio using a 10-point calibration curve constructed from a range of concentrations of the non-deuterated form of each analyte and a fixed amount of deuterated internal standard. Samples that did not fall in the linear portion of the curve were excluded from the analysis.

*FAAH rs324420 SNP (C385A) Genotyping*

Genotyping of the *FAAH* rs324420 SNP (C385A) was carried out on DNA extracted from the whole blood of study participants. DNA isolation was performed using the Nucleospin™ DNA Blood Isolation Kit (Machery Nagel, Fisher Scientific, Dublin, Ireland) according to the manufacturer’s instructions. DNA concentration, purity, and integrity were determined using a Nanodrop spectrophotometer (ND-1000; Nanodrop, Labtech International, UK). Purity was assessed by the absorbance ratio at 260/280, with values of ∼1.8 considered acceptable, while integrity was evaluated with 260/230 ratio limits set at 2.0-2.2. DNA was equalised to 1.7 ng/uL (for loading approximately 20ng/well) and stored at -20°C until use. A predesigned TaqMan primer for human FAAH (rs324420) [Assay ID: C1897306_10] was used with universal genotyping mastermix (Applied Biosystems, Fisher Scientific, Dublin, Ireland). Genotyping assays were performed according to the manufacturer’s protocol using an Applied Biosystems ‘StepOne Plus’ PCR machine (BioSciences, Dun Laoghaire, Ireland) and its proprietary allelic discrimination software. Samples were removed from the -80°C freezer and left to thaw at room temperature for two hours. All reagents and the standard curve were prepared as per manufacturer’s protocol. Samples were diluted with sample dilution buffer. The pre-coated 96-well plate was washed twice with wash buffer using a plate washer (Agilent BioTek ELx50, Dublin, Ireland). 50µl of the standards and samples were pipetted into the plate’s wells in duplicate, followed immediately with 50µl of Biotin-labelled antibody working solution into each well. The plate was then sealed and incubated at 37°C for 45 minutes. The plate was removed and washed three more times with wash buffer. 100µl of HRP-Streptavidin conjugate was added to each well, before the plate was covered and incubated at 37°C for 30 minutes. Following this, five further washes were completed. 90µl of TMB substrate was added to each well, then the plate was covered and incubated in the dark for 10 minutes, allowing a blue colour to develop. As soon as this colour developed, the plate was removed from the incubator and 50µl of stop solution was added to each well and mixed thoroughly, leading to a yellow colour change. The plate was again covered and immediately brought to the microplate reader (HIDEX Sense Microplate Reader, Tulku, Finland), where the absorbance was read at 450nm. A four-parameter logistic curve was then plotted to establish the concentration of the samples relative to the optical density (OD) of the known standard curve.

***Statistics***

**Placebo-induced Circulating Analyte Concentration Changes**

To examine whether circulating analyte concentrations changed as a function of condition (placebo vs. control), timepoint (pre vs. post), or their interaction, separate 2 × 2 repeated measures ANOVAs were conducted for each analyte in jamovi (version 2.6).

**Supplementary Results**

***Placebo-induced Circulating Analyte Concentration Changes***

There was no significant main effect of condition, timepoint, or condition-by-timepoint interaction on eCB levels (AEA, 2-AG), NAE levels (OEA, PEA) or β-endorphin levels (all p > 0.05, see Table S15). These results indicate that raw circulating concentrations of eCBs, NAEs, and β-endorphin did not differ significantly between conditions or across timepoints at the group level. Importantly, this does not preclude individual differences in analyte mobilization from being associated with placebo analgesia, as tested in the linear mixed models reported in the main text.

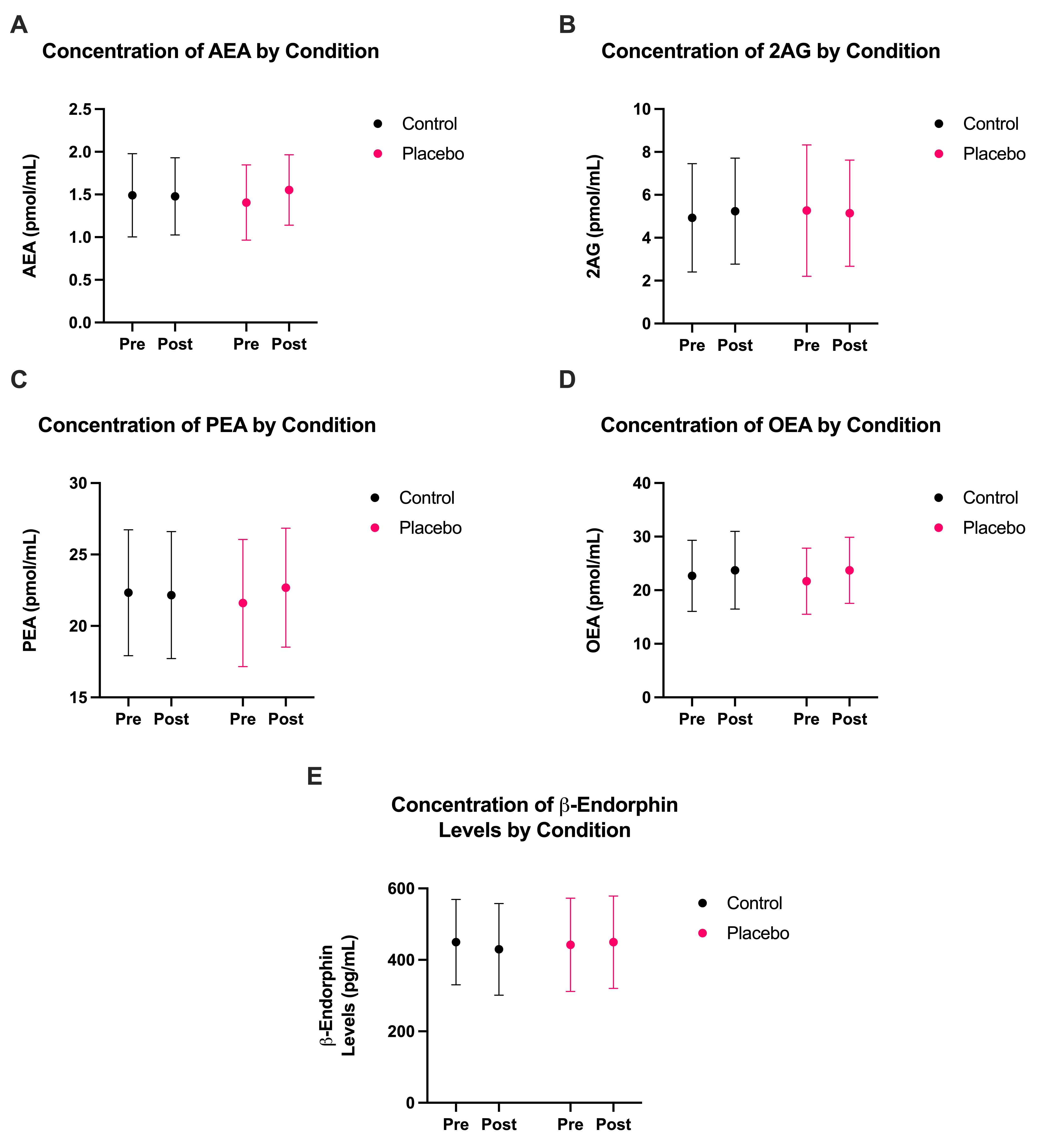

**Fig. S1. Circulating endocannabinoid, N-acylethanolamine, and β-endorphin concentrations before and after control and placebo conditions.** Mean pre- and post-condition concentrations are shown for control (black) and placebo (pink) conditions for (**A**) anandamide (AEA; pmol/mL), (**B**) 2-arachidonoylglycerol (2-AG; pmol/mL), (**C**) palmitoylethanolamide (PEA; pmol/mL), (**D**) oleoylethanolamide (OEA; pmol/mL), and (**E**) β-endorphin (pg/mL). Pre-condition values represent the blood draw immediately preceding each condition; post-condition values represent the blood draw immediately following. Error bars represent SD. N = 48. Repeated measures ANOVA revealed no significant interactions or main effects for any of the **endocannabinoid, N-acylethanolamine, and β-endorphin concentrations (all p > 0.05).**

**Table S9. Moderation of Placebo Analgesia by FAAH C385A Genotype, Cannabis Use, and Biological Sex.**

| **Effect** | | ***F*** | | ***df*** | ***p*** | | **Partial *η*²** |
| --- | --- | --- | --- | --- | --- | --- | --- |
| ***FAAH C385A Genotype*** | | | | | | | |
| Condition × Genotype interaction | | 0.78 | | (1, 46) | .381 | | .017 |
| Main effect of genotype | | 3.35 | | (1, 46) | .074 | | .068 |
| ***Cannabis Use History*** | | | | | | | |
| Condition × Cannabis Use interaction | | 0.26 | | (1, 46) | .610 | | .006 |
| Main effect of cannabis use | | 3.16 | | (1, 46) | .082 | | .064 |
| ***Biological Sex*** | | | | | | | |
| Condition × Sex interaction | | 2.84 | | (1, 46) | .099 | | .058 |
| ***Estimated Marginal Means — Biological Sex*** | | | | | | | |
|  | **Mean (Control) ± SEM** | | **Mean (Placebo) ± SEM** | | | **Δ (Placebo − Control)** | |
| Females | 3.57 ± 0.27 | | 2.19 ± 0.33 | | | −1.38 | |
| Males | 3.37 ± 0.26 | | 2.62 ± 0.32 | | | −0.75 | |

**Note.** All tests used mixed-measures ANOVA with condition (placebo vs. control) as a within-participants factor. *F* = *F*-statistic; *df* = degrees of freedom; *p* = *p*-value; partial *η*² = partial eta-squared effect size. SEM = standard error of the mean; Δ = difference.

Table S10. Spearman correlations among baseline FAAH substrate levels.

|  | **AEA** | **PEA** | **OEA** |
| --- | --- | --- | --- |
| **AEA** | — |  |  |
| **PEA** | 0.749*** | — |  |
| **OEA** | 0.735*** | 0.759*** | — |

Note. N = 48. AEA = anandamide; PEA = palmitoylethanolamide; OEA = oleoylethanolamide. ***p < .001.

Table S11. Spearman correlations among FAAH substrate changes during control.

|  | **AEA** | **PEA** | **OEA** |
| --- | --- | --- | --- |
| **AEA** | — |  |  |
| **PEA** | 0.646*** | — |  |
| **OEA** | 0.791*** | 0.789*** | — |

**Note.** N = 48. AEA = anandamide; PEA = palmitoylethanolamide; OEA = oleoylethanolamide. ***p < .001.

Table S12. Spearman correlations among FAAH substrate changes during placebo.

|  | **AEA** | **PEA** | **OEA** |
| --- | --- | --- | --- |
| **AEA** | — |  |  |
| **PEA** | 0.668*** | — |  |
| **OEA** | 0.821*** | 0.768*** | — |

**Note.** N = 48. AEA = anandamide; PEA = palmitoylethanolamide; OEA = oleoylethanolamide. ***p < .001.

Table S13. Principal component analysis assumption tests comparing inclusion versus exclusion of 2-AG in FAAH substrates component.

| **Condition** | **Variables** | **Bartlett's χ²** | **df** | **p** | **KMO** | **2-AG KMO** | **Eigenvalue** | **% Var** | **2-AG Load** |
| --- | --- | --- | --- | --- | --- | --- | --- | --- | --- |
| Baseline | AEA, 2-AG, PEA, OEA | 108.265 | 6 | <0.001 | 0.695 | 0.314 | 2.668 | 65.81 | 0.178 |
| Baseline | AEA, PEA, OEA | 99.295 | 3 | <0.001 | 0.757 | — | 2.615 | 86.05 | — |
| Control | ΔAEA, Δ2-AG, ΔPEA, ΔOEA | 65.880 | 6 | <0.001 | 0.704 | 0.775 | 1.523 | 65.30 | 0.176 |
| Control | ΔAEA, ΔPEA, ΔOEA | 59.820 | 3 | <0.001 | 0.679 | — | 1.486 | 75.45 | — |
| Placebo | ΔAEA, Δ2-AG, ΔPEA, ΔOEA | 110.310 | 6 | <0.001 | 0.738 | 0.812 | 1.258 | 65.97 | 0.442 |
| Placebo | ΔAEA, ΔPEA, ΔOEA | 100.365 | 3 | <0.001 | 0.715 | — | 1.100 | 84.61 | — |

**Note.** Bartlett's test of sphericity assessed whether the correlation matrix differed significantly from an identity matrix (p < 0.05 indicates suitability for PCA). Kaiser-Meyer-Olkin (KMO) measure of sampling adequacy assessed the proportion of variance that might be common variance (values > 0.5 indicate suitability for PCA). 2-AG KMO = individual KMO value for 2-AG; 2-AG Load = PC1 loading for 2-AG. When 2-AG was included, it failed to meet criteria at baseline (KMO = 0.314) and showed low PC1 loadings across conditions (0.18–0.44), indicating it does not share common variance with FAAH substrates. This is consistent with 2-AG being primarily metabolized by monoacylglycerol lipase rather than FAAH. The final FAAH-substrates component therefore includes only AEA, PEA, and OEA, with 2-AG analyzed separately. Δ = change scores (post-condition minus pre-condition). N = 48 participants.

**Table S15. ANOVA Table for Placebo-Induced Analyte Changes**

|  | ***F*** | ***df*** | ***p*** |
| --- | --- | --- | --- |
| ***AEA*** |  |  |  |
| Condition | 0.005 | 1 ,47 | 0.943 |
| Timepoint | 1.063 | 1 ,47 | 0.308 |
| Condition × Timepoint | 2.495 | 1 ,47 | 0.121 |
| ***2-AG*** |  |  |  |
| Condition | 0.116 | 1 ,47 | 0.735 |
| Timepoint | 0.053 | 1 ,47 | 0.819 |
| Condition × Timepoint | 0.506 | 1 ,47 | 0.481 |
| ***OEA*** |  |  |  |
| Condition | 0.445 | 1 ,47 | 0.508 |
| Timepoint | 2.505 | 1 ,47 | 0.12 |
| Condition × Timepoint | 0.388 | 1 ,47 | 0.536 |
| ***PEA*** |  |  |  |
| Condition | 0.033 | 1 ,47 | 0.857 |
| Timepoint | 0.397 | 1 ,47 | 0.532 |
| Condition × Timepoint | 1.645 | 1 ,47 | 0.206 |
| ***β-endorphin*** |  |  |  |
| Condition | 0.254 | 1 ,47 | 0.616 |
| Timepoint | 0.384 | 1 ,47 | 0.539 |
| Condition × Timepoint | 1.068 | 1 ,47 | 0.307 |

**Supplementary References**

1.	Amir C, Rose-McCandlish M, Weger R, Dildine TC, Mischkowski D, Necka EA, et al. (2022): Test-Retest Reliability of an Adaptive Thermal Pain Calibration Procedure in Healthy Volunteers. *J Pain*. 23:1543-1555.
